# Muscle Metabolic Energy Costs While Modifying Propulsive Force Generation During Walking

**DOI:** 10.1101/2020.07.31.230698

**Authors:** Richard E. Pimentel, Noah L. Pieper, William H. Clark, Jason R. Franz

## Abstract

We pose that an age-related increase in the metabolic cost of walking arises in part from a redistribution of joint power where muscles spanning the hip compensate for insufficient ankle push-off and smaller peak propulsive forces (F_P_). Young adults elicit a similar redistribution when walking with smaller F_P_ via biofeedback. We used targeted F_P_ biofeedback and musculoskeletal models to estimate the metabolic costs of operating lower limb muscles in young adults walking across a range of F_P_. Our simulations support the theory of distal-to-proximal redistribution of joint power as a determinant of increased metabolic cost in older adults during walking.

## Introduction

Aging and many gait pathologies are associated with reduced walking economy (i.e., higher metabolic cost when walking at the same speed). People who require greater metabolic cost to walk fatigue more quickly and are thereby likely to walk less. The functional consequences of higher metabolic cost include reduced physical activity, independence, and thereby quality of life (Morris & Hardman 1997). Moreover, this barrier to accessible exercise disproportionally impacts populations already at increased risk for secondary health concerns such as cardiovascular disease and diabetes (World Health Organization 2019). Preserving and/or restoring walking ability can be transformative in improving physical activity and overall health (Morris & Hardman 1997). Identifying the mechanistic determinants of higher metabolic cost of walking due to age and gait pathology is a critical first step toward translational opportunities to preserve and/or restore economical walking patterns in people with walking disability.

The higher metabolic cost of walking due to aging, in particular, is likely to affect a significant and ever-increasing proportion of our population. Older adults consume oxygen 20-30% faster than younger adults while walking at the same speed (Peterson & Martin 2010; Hortobágyi et al. 2011). The reasons for this higher metabolic energy cost are not fully understood. Training interventions shown to improve strength and/or balance in older adults have been generally unable to reduce the metabolic cost of walking (Mian et al. 2007; Franz 2016). Several plausible explanations for this increased metabolic cost of walking have emerged and align with the neurological, muscular, and physiological changes due to aging. Some explanations, including higher metabolic costs to maintain lateral stability or to perform more total mechanical work during walking, have been refuted (Ortega & Farley 2007; Ortega et al. 2008). Others, such as increased antagonist leg muscle coactivation in older adults, leave roughly two-thirds of the age-related increase in metabolic cost unexplained (Peterson & Martin 2010; Hortobágyi et al. 2011).

Biomechanically, older adults walk with a hallmark redistribution of muscular workload wherein muscles spanning the hip during early stance (i.e., hip extensors) and/or late stance and early swing (i.e., hip flexors) increase their output presumably to compensate for reduced ankle push-off – so described as a distal-to-proximal redistribution of muscular demand (Kerrigan et al. 1998; DeVita & Hortobagyi 2000; Silder et al. 2008; Cofré et al. 2011; Boyer et al. 2017). At the limb-level, this reduced ankle push-off in older adults is accompanied by smaller peak propulsive ground reaction forces (F_P_) – an important determinant of walking speed (Boyer et al. 2017). This likely leads to a reduction in their preferred walking speed, which is a proxy for functional ability, mortality, and cognitive decline, with many health experts suggesting that walking speed may be a vital sign of health status of older adults (Hardy et al. 2007; Kikkert et al. 2016; Jee et al. 2019). Growing indirect evidence suggests that a distal-to-proximal redistribution of muscular workload during walking may come with a significant metabolic penalty rooted in inter-muscular differences in muscle-tendon architecture. Specifically, with their short muscle fibers and long, compliant series elastic tendons, we (Fickey et al. 2018; Browne & Franz 2019) and others (Sawicki et al. 2009; Huang et al. 2015) have suggested that muscle-tendon units spanning the ankle are far more economical for force generation in walking than those spanning the hip, which have much longer muscle fibers and relatively shorter and less compliant tendons. However, despite its potential relevance to walking economy in aging and gait pathology, this theory has not yet been supported by direct experimental manipulation or measurement.

It is challenging to experimentally isolate hallmark features of elderly gait such as the distal-to-proximal redistribution and/or reduced F_P_. Recently, our group showed that young adults walking with smaller than normal F_P_ in a targeted biofeedback paradigm redistribute muscle workload from the ankle to the hip without changing total positive mechanical work, thereby emulating the walking biomechanics of older adults (Browne & Franz 2017a; Fickey et al. 2018). Thus, using this biofeedback paradigm in young adults would allow for testing how differences in the relative reliance on distal versus proximal leg muscles that typically accompany reductions in F_P_ affect muscle metabolic cost. Additionally, a young adult demographic allows for biomechanical investigation that is independent of other age-related changes that may affect metabolism (sarcopenia, tendon compliance, mitochondrial physiology, etc.).

It is not feasible to directly measure the metabolic cost of operating individual muscles during walking. Fortunately, models of the human musculoskeletal system allow for simulation of body movements and estimation of muscle dynamics underlying those movements. Thus, our purpose was to use a targeted biofeedback paradigm and musculoskeletal simulations to estimate the metabolic costs of operating lower limb muscles in young adults walking across a range of F_P_. We first hypothesized that the simulated muscle metabolic energy cost of walking would increase in a manner consistent with whole-body measurements when targeting smaller or larger F_P_. We also hypothesized that the metabolic costs of operating proximal versus distal leg muscles would exhibit fundamentally different responses to targeting smaller and larger F_P_. Ultimately, we seek to provide muscle-level insight to inform movement scientists, rehabilitation engineers, and clinicians interested in understanding and mitigating age- and disease-related increases in walking metabolic cost.

## Methods

### Participants

We recruited 12 young adults (7 females; age=23.3±3.14 years; height=1.74±0.12 m; mass=74.7±14.3 kg, BMI=24.6±3.0 kg/m^2^, *mean ± standard deviation*) via word of mouth to participate in this single-visit study. All participants confirmed absence of neurologic impairments and musculoskeletal injuries in the previous 6 months. Prior to data collection, each participant provided written informed consent and the study was approved by the University of North Carolina Biomedical Sciences Institutional Review Board.

### Experimental Protocol

We measured each participant’s preferred over-ground walking speed via the average time from three passes down a 30-meter walkway (average preferred walking speed: 1.37±0.15 m/s). For motion capture measurements, we placed 19 retroreflective markers on the sacrum and bilateral anterior superior iliac spines, posterior superior iliac spines, lateral/medial femoral epicondyles, lateral/medial malleoli, lateral calcanei, and lateral first and fifth metatarsal heads. An additional 14 markers tracked the thighs and shanks of each participant on rigid clusters. After a static calibration trial, we located functional hip joint centers via leg circumduction tasks to improve model scaling accuracy (Cappozzo 1984; Bell et al. 1990; Piazza et al. 2001). Prior to walking measurements, participants stood still for 5 minutes to estimate standing metabolic rate (see *Metabolic Measurements*). Participants then completed all walking trials at their preferred speed on a force-sensing, dual-belt treadmill (Bertec Corp., Columbus, Ohio, USA), beginning with a 5-minute warm up. The average peak anterior ground reaction force from all steps taken during the final two minutes of the warm-up determined each subjects’ habitual F_P_ using a custom Matlab (MathWorks, Natick, MA, USA) script that extracted and analyzed ground reaction forces in real-time.

Subjects then participated in a visual biofeedback paradigm in which they were encouraged to push off the ground with varying vigor to target their typical walking F_P_ (Norm) as well as ±20% and ±40% of Norm as outlined below (Figure 1A). Using a Matlab script and visual interface described in detail previously (Browne & Franz 2017a), a monitor positioned in front of the participant simultaneously displayed the respective target F_P_ value for each trial and the average F_P_ from their previous 4 steps (2 right and 2 left), updated on a step-by-step basis (Browne & Franz 2017a). For all trials involving biofeedback, we normalized the scaling of each subject’s F_P_ data on the monitor to evenly distribute all target values over the ordinate range. Prior to beginning the targeting biofeedback, each participant completed an exploration period to become familiarized with increasing and decreasing their F_P_ across the range of target values. Then, across five separate 5-minute trials, participants walked at their preferred speed and targeted each of the F_P_ targets in randomized order. During the final two minutes of each trial, we recorded bilateral lower body kinematics using a 12-camera motion capture system (Motion Analysis Corporation, Santa Rosa, CA, USA) operating at 100 Hz and measured split-belt treadmill ground reaction forces at 1000 Hz. We filtered marker trajectories and ground reaction forces using a 4^th^-order low-pass Butterworth filter with 6 and 20 Hz cutoff frequencies, respectively.

**Figure 1:**
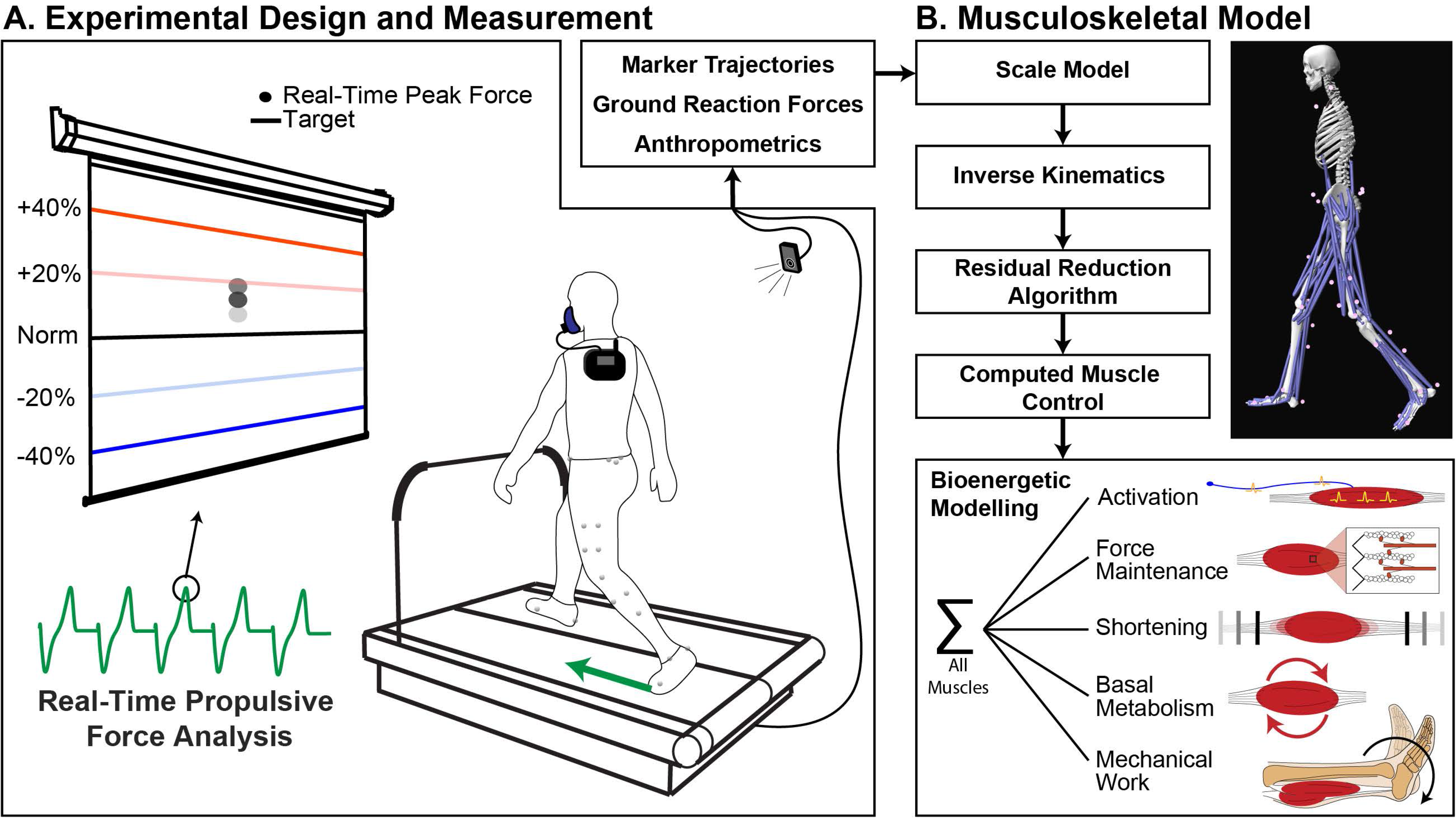
Participants pushed off the ground with varying vigor to target −40%, −20%, +20%, +40%, and their typical (Norm) propulsive force (F_P_) in a walking biofeedback paradigm with 5-minute trials per condition. Participant anthropometrics, marker trajectories, and ground reaction forces from the final 2 minutes of each trial defined musculoskeletal modeling simulations. We simulated the first left and right full stride from the 10 second window with the most accurate biofeedback targeting performance. From these simulations, we ultimately determined the metabolic power required from each lower body muscle to drive the scaled model and match the measured movements.

### Metabolic Measurements

For this study, we repurposed metabolic measurements from a previously published study using methods therein described in detail (Pieper et al. In Revision). Briefly, expired oxygen and carbon dioxide were sampled on a breath-by-breath basis via COSMED K5 wearable metabolic system (COSMED, Rome, Italy). To estimate standing and walking metabolic rates, we averaged expired gases over the final two minutes of the 5-minute standing condition and each 5-minute walking condition, respectively. We used standardized regression equations (Brockway 1987) to estimate whole-body metabolic power from oxygen consumption and carbon dioxide production, subtracted standing from walking to calculate net metabolic power, and finally normalized outcomes by body mass (W/kg).

### Inclusion of Trunk Kinematics

Our original walking data was derived from another study (Pieper et al. In Revision) and did not include trunk marker trajectories; however, the reliability of musculoskeletal simulations improves when including trunk kinematics for center of mass estimates and dynamic consistency. Thus, we opted to record and benchmark normal pelvis and trunk marker trajectories (adding sternal notch and bilateral acromion processes) in three additional participants walking at 1.2 m/s. We then used the resultant average kinematics to create a virtual trunk segment for each participant in our primary experiment. Specifically, we synchronized the measurements using time-normalized walking motions and defined virtual trunk marker locations as position vectors in the pelvis coordinate system. We scaled the position vector magnitudes to each participant’s average leg length to account for inter-individual differences in subject stature. Supplementary Figure S1 shows the profiles of virtual trunk kinematics across all subjects and conditions.

### Musculoskeletal Modeling and Simulation

We performed all musculoskeletal modeling and bioenergetic simulations using filtered marker trajectories and ground reaction forces in OpenSim (version 4.0) (Delp et al. 2007; Seth et al. 2018) and the gait2392 musculoskeletal model (Figure 1B). We preserved all default muscle parameters, except maximum isometric force, which was set to 1.5 times default for all models (in accordance with OpenSim documentation, and similarly performed by other groups (Arnold et al. 2013)). Subject-specific models were scaled using a combination of anatomical, functional, and virtual markers that aligned with the local coordinate axes of each body segment. We locked the subtalar and metatarsal-phalangeal joints at 0° for all simulations because our marker set was insufficient to yield those kinematics (Rajagopal et al. 2016).

We analyzed the first left and right strides from the 10-second window with the most accurate biofeedback performance (i.e., steps where their actual F_P_ closely matched the target) for each walking condition and subject. From these two strides, inverse kinematics determined segment kinematics that minimized the sum of the square marker trajectory error. Then, we iteratively applied a residual reduction algorithm to identify model segment mass distributions and joint kinematics consistent with measured ground reaction forces. We updated model segment masses after the first iteration and used kinematics from the second iteration to ensure dynamic consistency. After residual reductions, we used computed muscle control without constraints to determine the lower limb muscle activations necessary to drive the scaled model to match the recorded kinematics.

Finally, we determined our primary outcome measure (muscle metabolic power) using two different muscle metabolic cost models (i.e., “probes”) readily available within OpenSim (Umberger et al. 2003; Bhargava et al. 2004; Umberger 2010). Our purpose was not to objectively compare the models’ efficacy; rather, we included two models with checks for consistency to build confidence in our model predictions. Each model estimated metabolic power for each individual muscle actuator as a function of heat rate (activation, shortening/lengthening, force maintenance, basal) and mechanical work rate (Umberger et al. 2003; Bhargava et al. 2004; Umberger 2010). It is not within the scope of this paper to discuss these models in detail. Nevertheless, each was governed by time-dependent variables that included, for each individual muscle actuator, its excitation and activation, relative % of fast versus slow twitch muscle fibers, contractile velocity, force-generating capacity, mass, and optimal fiber length.

### Data Reduction and Statistical Analysis

We integrated individual muscle metabolic powers over select phases of the gait cycle (i.e., leading limb double support, single support, trailing limb double support, swing phase, and whole stride) which were calculated from instances where the vertical ground reaction force crossed a threshold of 20 N. In addition, we normalized all metabolic outcomes by body mass and total stride time to estimate metabolic power per unit second of walking. We categorized joint-level metabolic powers as those associated with operating the hip, knee, and ankle joints during each gait cycle phase by summing individual muscle metabolic powers for actuators spanning each respective joint. We partitioned biarticular muscles into their respective joints using the ratio of their sagittal plane moment arms in a standing position (Miller 2014). Finally, we averaged individual muscle metabolic outcomes bilaterally for subsequent statistical analysis.

Pearson correlations tested for associations between measured and model-predicted metabolic powers. One-way repeated measures analyses of variance (ANOVA) tested for main effects of biofeedback condition on joint-level metabolic powers within each gait cycle phase of interest. When a significant main effect was found, Fisher’s least significant difference (LSD) post-hoc tests identified pairwise differences compared to Norm. One-dimensional statistical parametric mapping (SPM, http://www.spm1d.org/, (Pataky 2012)) identified regions of the gait cycle over which individual muscle metabolic powers during targeted biofeedback conditions differed from Norm (repeated measures ANOVA with LSD post-hoc tests). We performed correlations and ANOVAs in Matlab, and calculated SPM using Python. We used a critical alpha value of 0.05 to define significance of ANOVAs. We additionally report effect sizes of partial eta square (*ղ*_p_^2^) for the variance associated with main effect across biofeedback conditions as well as Cohen’s d for effect size pairwise comparisons between a specific condition versus Norm.

## Results

### Empirical Measurements versus Model Predictions

Compared to walking normally, measured net metabolic power increased by an average of 58% and 47% when targeting −40% and +40% F_P_, respectively (Figure 2A, p<0.001, *ղ*_p_^2^=0.350). Total model-predicted metabolic power followed a similar trend (Figure 2B-C) and positively correlated with empirical measurements (Umberger: r^2^=0.309, p<0.001; Bhargava: r^2^=0.342, p<0.001, Figure 2E-F). These two metabolic models were strongly correlated with each other (r^2^=0.981, p<0.001, Figure 2D) and displayed similar stride-averaged profiles (Figure 2G). Given the high level of consistency between model predictions (Figure 2E, F, G), we opted to use the Umberger model for subsequent joint- and muscle-level analysis.

**Figure 2:**
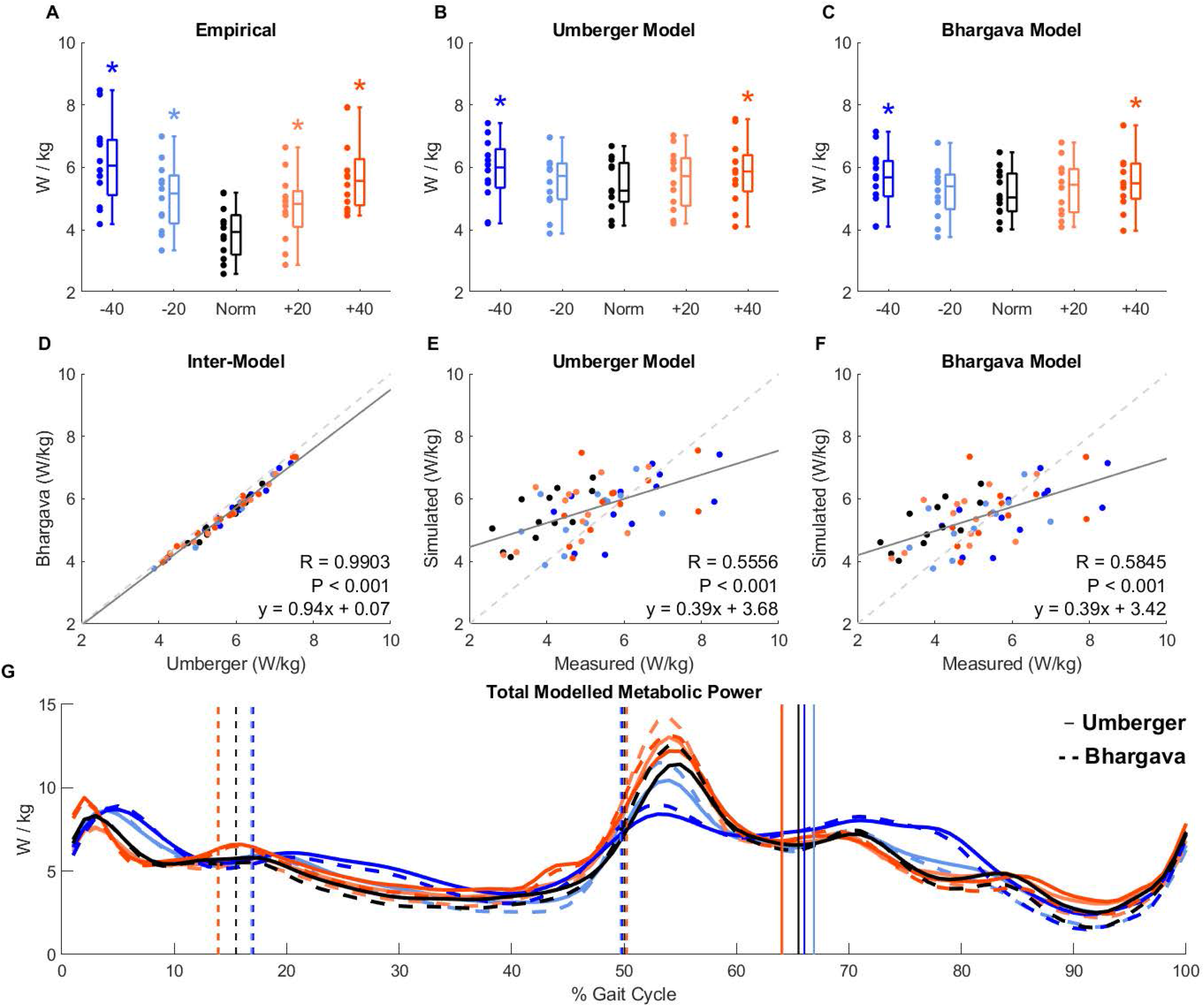
For each biofeedback condition, we display the walking metabolic power empirically measured using indirect calorimetry (A) and simulated from two different musculoskeletal metabolic models (B, C). The models produced nearly identical results (D) that correlated moderately with empirically-measured metabolic power (E, F), explaining about 1/3^rd^ of the variance. Musculoskeletal modeling of metabolic power allows researchers to estimate the metabolic power required for lower body muscles to drive a simulation of each participant’s individual walking pattern. This allows for comparison across the F_P_ biofeedback conditions at specific instances of the gait cycle (G). Given the high similarity of the two models, we opted to use the Umberger model for our metabolic cost estimates and statistical comparisons.

### Joint-Level Model Predictions

Figure 3 summarizes model-predicted metabolic power integrated across periods of interest for all muscles (A) and summed for muscle actuators spanning the hip (B), knee (C), and ankle (D) joints. Due to the many statistical comparisons for the joint-level analysis, please reference Table 1 where we document the statistical outcomes of all comparisons. Here, we report only mean pairwise differences for brevity. When targeting 40% smaller F_P_, metabolic power across the entire stride increased in total (+0.79 W/kg) due to increases at the hip (+0.29 W/kg) and knee (+0.22 W/kg). These increases were mainly attributed to increased power consumed during leading leg double support at the hip (+0.16 W/kg) and during leg swing in total (+0.42 W/kg) and at the knee (+0.11 W/kg). Also, when targeting 40% smaller F_P_, ankle metabolic power increased during single support (+0.07 W/kg) but decreased during trailing limb double support (−0.15 W/kg), resulting in no change over the entire stride.

**Table 1:**
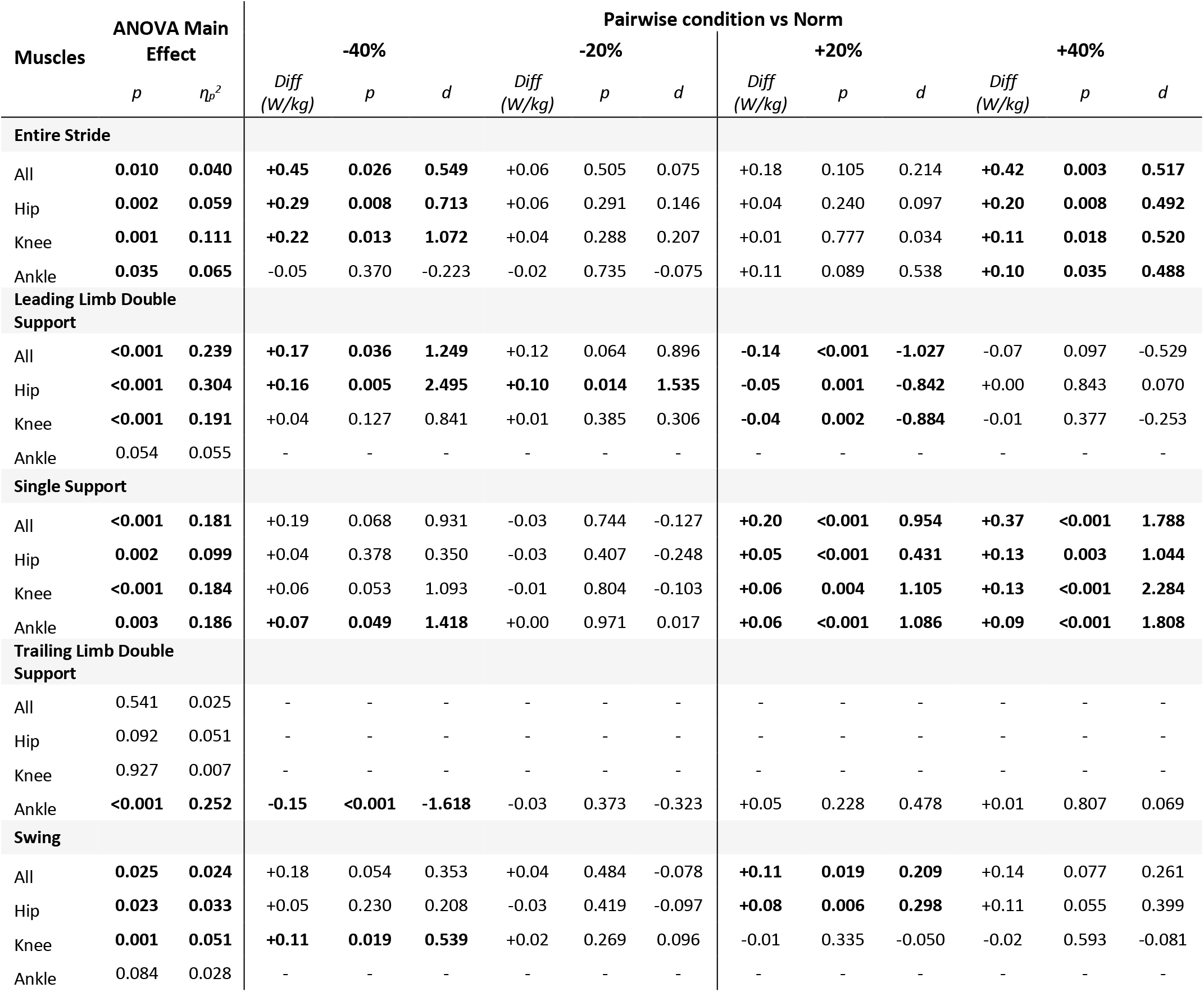
Statistics for repeated measures ANOVA and LSD post-hoc pairwise tests corresponding to Figure 3. Conditions with significant p-values are bolded. We do not report pairwise differences for joint-level metabolic costs with insignificant ANOVA main effect (rows with missing data). ղ_p_^2^: partial eta squared, *d*: Cohen’s D, *p*: pairwise p-value, Diff: mean difference versus Norm.

**Figure 3:**
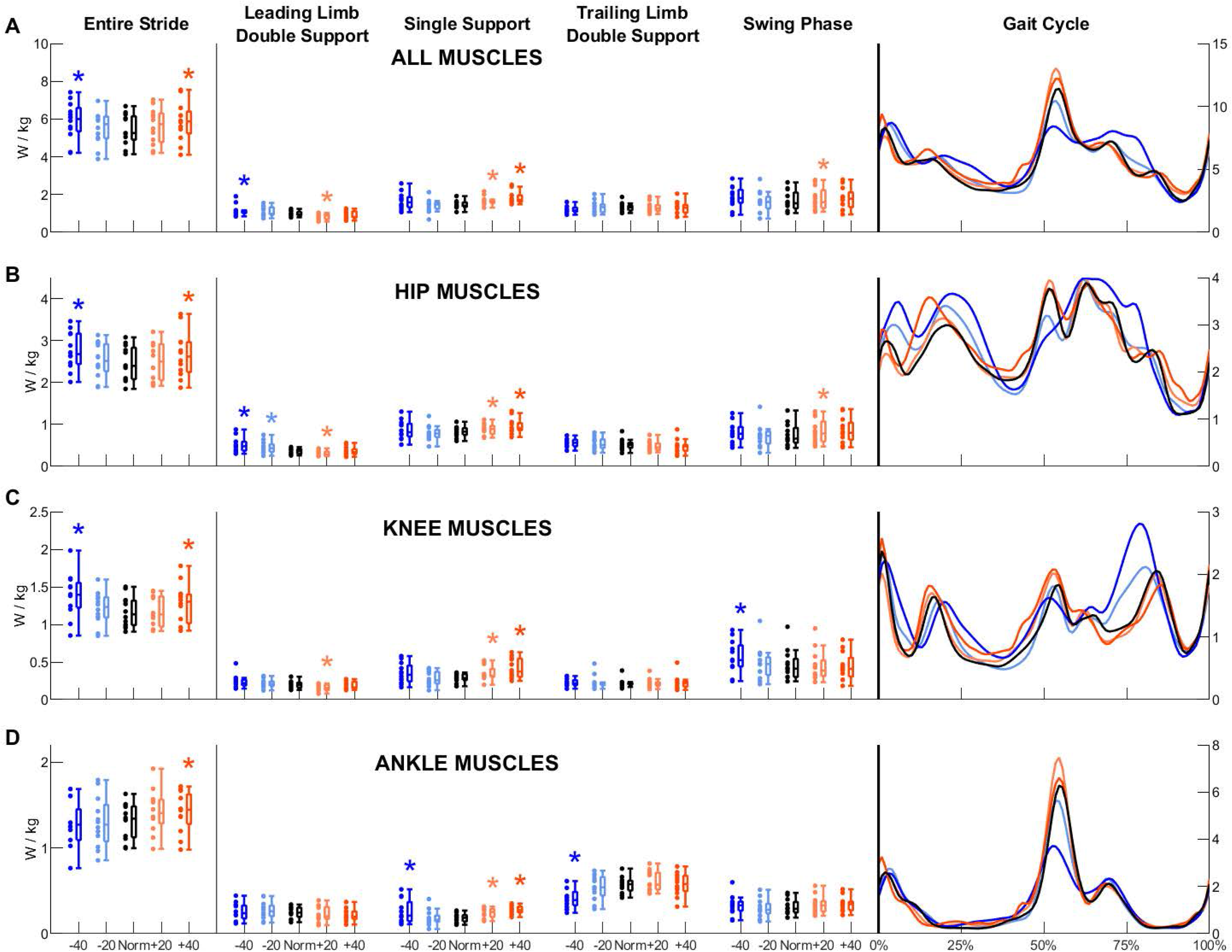
For each biofeedback condition, we show net metabolic power summed for all modeled lower body muscles (A), and for those spanning each of the major joints of the lower limb (B, C, D). Metabolic powers were integrated over: the entire stride, leading limb double support, single limb support, trailing limb double support, and swing phase. Asterisks show significant LSD post hoc pairwise differences versus Norm in the presence of a significant ANOVA main effect. On the right, we also display instantaneous net metabolic power across the gait cycle.

When targeting 40% larger F_P_, metabolic power across the entire stride increased in total (+0.90 W/kg) due to increases at the hip (+0.20 W/kg), knee (+0.11 W/kg), and ankle (+0.10 W/kg). These increases over the entire stride were primarily due to higher metabolic power during single support across all lower limb joints (hip: +0.13 W/kg, knee: +0.13 W/kg, ankle: +0.09 W/kg). We found no differences in metabolic power during swing phase when targeting 40% larger F_P_. However, when targeting 20% larger F_P_, swing phase metabolic power increased in total (+0.11 W/kg) and at the hip (+0.08 W/kg). Finally, there was a reduction in metabolic cost during leading limb double support when targeting +20% F_P_ in total (−0.14 W/kg) and at the hip (−0.05 W/kg) and knee (−0.04 W/kg), which were the only significant reductions in metabolic cost across all conditions.

### Individual Muscle-Level Model Predictions

Figure 4 shows the metabolic power estimated for 21 lower body muscles that contribute most to total model-predicted metabolic power during walking and the results of our statistical parametric mapping. Although there were several instances containing within-subject condition main effects, only a handful of significant pairwise differences arose and nearly all were for the 40% smaller F_P_ condition. Specifically, when targeting 40% smaller F_P_, we observed the following changes in metabolic power (organized from distal to proximal): soleus consumed less during push-off; biceps femoris (long head) and semimembranosus consumed more during early single support; the vastii muscles consumed less during weight acceptance and more during mid-swing; and the gluteus medius and maximus both consumed less during push-off and more during late swing/early stance. When targeting 20% smaller F_P_, we found that the vastus medialis and intermedius consumed more metabolic power during mid-swing. Finally, when targeting 40% larger F_P_, the adductor longus consumed less metabolic power during early single support.

**Figure 4:**
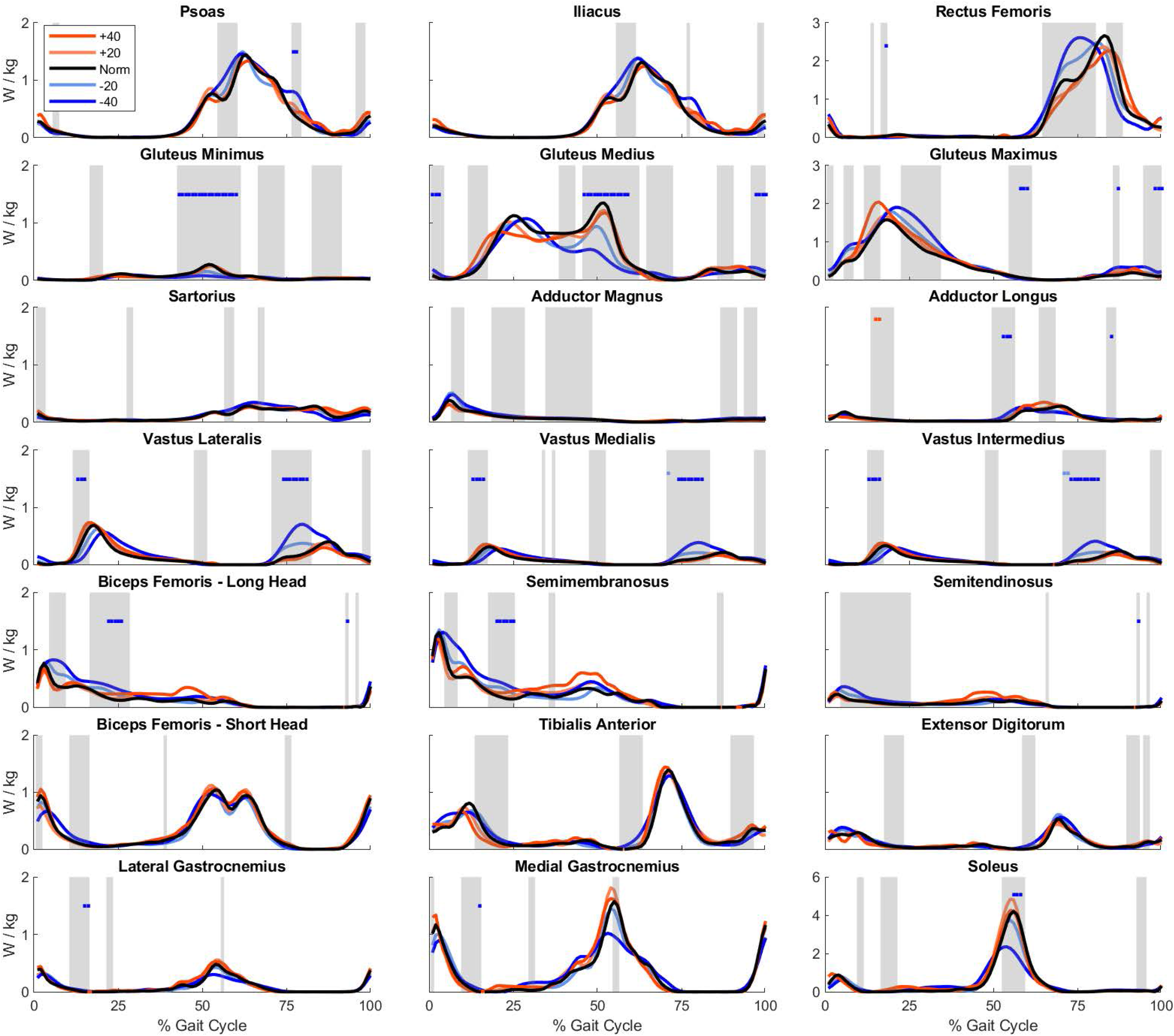
We plot the metabolic powers across the gait cycle for the 21 highest power-consuming muscles of the lower limb for each biofeedback condition. Gray shaded regions show periods with a significant repeated measures ANOVA main effect for biofeedback condition using statistical parametric mapping. Colored blocks identify any LSD post hoc pairwise differences for that respective colored condition versus Norm. OpenSim muscles with multiple lines of action (gluteus minimus/medius/maximus and adductor magnus) were summed for simplicity.

### Step Kinematics

To provide context for our metabolic outcomes, Table 2 displays timing information for gait cycle periods of interest. Participants took longer (shorter) steps when target larger (smaller) than normal F_P_, respectively. In relative terms (i.e., % stride), participants spent more time in single support and leg swing and less time in double support when targeting larger F_P_.

**Table 2:**
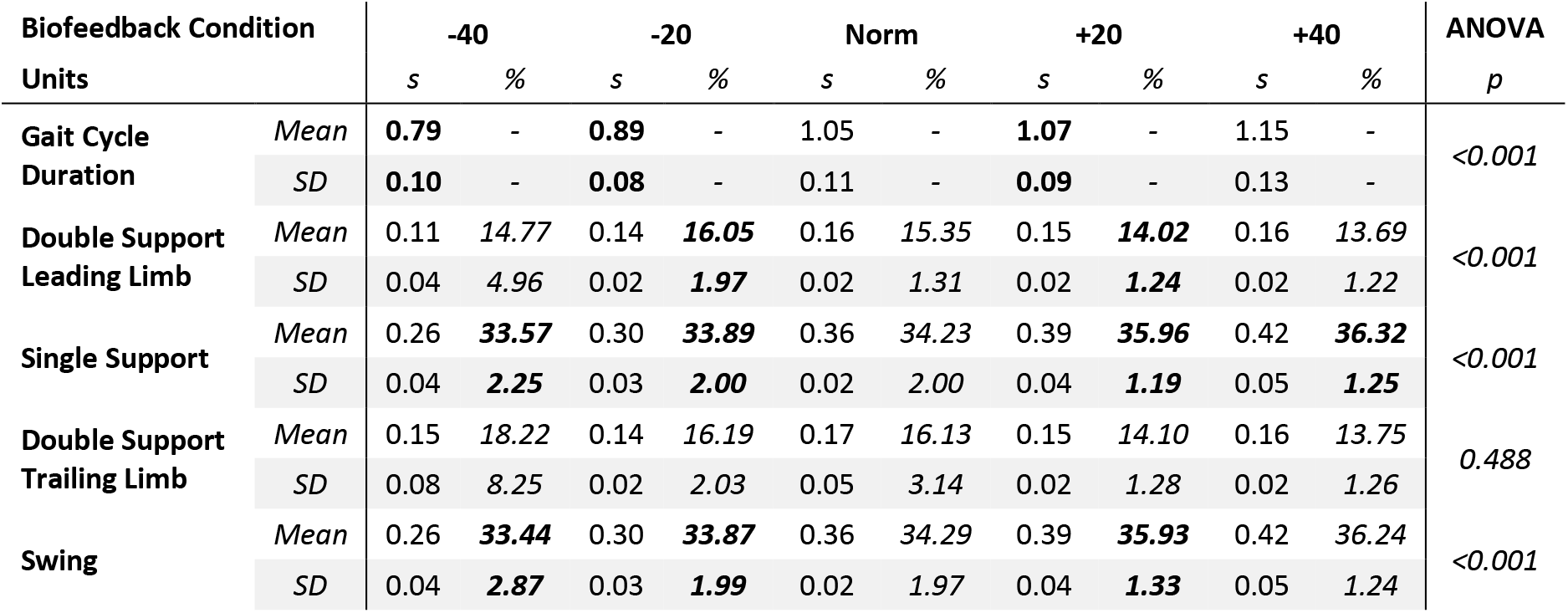
Gait cycle periods of interest for each biofeedback condition in absolute (s) and relative (%) time. These periods were calculated over 10 strides in the window of best biofeedback targeting performance (i.e., best matching the −40%, −20%, etc. of F_P_). Bolded values indicate significant post-hoc pairwise difference versus Norm. These are the periods over which the joint-level metabolic costs were integrated in Figure 3. Stride time is defined as 100% of the gait cycle, thus we omit these data and run post hoc analyses using duration. SD: standard deviation across participant means, s: seconds, %: percent of gait cycle

## Discussion

This study sought to provide new insight into the biomechanical basis for reduced walking economy due to age and gait pathology, and thereby inform movement scientists, rehabilitation engineers, and clinicians interested in mitigating those changes to improve independence. We accept our first hypothesis that model-predicted metabolic costs increased in a manner consistent with those measured across conditions via indirect calorimetry. We also accept our second hypothesis that proximal and distal leg muscles would exhibit different responses when targeting smaller and larger than normal F_P._ Ultimately, our experimental manipulations and musculoskeletal simulations provide a muscle-level roadmap for better understanding the metabolic consequences of walking with: (*i*) smaller than normal F_P_, common in aging and gait pathology, and (*ii*) larger than normal F_P_, a therapeutic milestone for gait retraining or wearable assistive devices. More immediately, our results provide convincing evidence to suggest that a distal-to-proximal redistribution of muscle workload during walking may exact a significant metabolic penalty.

In their seminal papers, Bhargava et al. (2004) reported model predictions that tended to overestimate measured net metabolic power, while Umberger et al. (2003) and Umberger (2010) yielded moderately better agreement (Umberger et al. 2003; Bhargava et al. 2004). More recently, Koelewijn et al. (2019) found very strong positive correlations between the output from both these models and measured net metabolic power during walking (r^2^ = 0.92 and 0.86, respectively) (Koelewijn et al. 2019). In this study, our model predictions explained approximately one-third of the variance in empirical measurements across a relatively broad combination of walking conditions. When averaged across conditions, our estimates agreed quite well with measured values in absolute terms. However, we note two explanations for the variance unaccounted for across conditions. Specifically, our models tended to overestimate net metabolic power during normal walking while underestimating the sensitivity therein to prescribed changes in F_P_. Moving forward, we anticipate refining our predictions based on higher fidelity measurements (i.e., including trunk, subtalar, and metatarsal-phalangeal joint kinematics) and improved simulation specificity (i.e., personalized “tuning” of muscle-tendon architecture (Handsfield et al. 2014) and material properties (Uchida et al. 2016; Orselli et al. 2017)). Nevertheless, because we were able to estimate changes in whole-body metabolic cost consistent with those measured, we have confidence in using musculoskeletal modeling to answer questions related to the metabolic consequences of altered muscular demand across the lower limbs during walking.

As context for our discussion of metabolic outcomes, we do note variations in step kinematics between F_P_ conditions (Table 2), with shorter and faster steps when targeting smaller F_P_ and vice versa. Such changes in step kinematics are known to affect metabolic cost (Donelan et al. 2002; Kuo et al. 2005; Soo & Donelan 2010). Because the purpose of our study was to assess the metabolic costs of walking while controlling for speed, we cannot rule out the interaction between step length/duration and walking dynamics that govern our model estimates. However, we also know from our prior work that although both change simultaneously in response to our biofeedback paradigm, modifying F_P_ is not biomechanically equivalent to modifying step lengths (Browne & Franz 2017a). Hereafter, we consider metabolic costs at the joint- and muscle-level foremost as those due to changes in F_P_, but also in the context of these changes in gait event timing.

### Targeting smaller than normal F_P_

Reduced ankle power generation during push-off, characteristic of aging and many gait pathologies and associated with reduced F_P_, requires compensatory increases in hip mechanical output. In prior studies, this compensation has manifested as an increased demand on leading limb hip extensors to accelerate the body forward and upward (DeVita & Hortobagyi 2000; Silder et al. 2008; Cofré et al. 2011), and on trailing limb hip flexors to initiate leg swing (Judge et al. 1996; Silder et al. 2008; Cofré et al. 2011). Our results provide evidence that such compensations have metabolic consequences. First, as expected for walking with reduced push-off intensity, we found that targeting smaller F_P_ yielded lower metabolic costs of operating muscles spanning the ankle during trailing limb double support. More proximally, we found that increases in the metabolic cost of walking with smaller F_P_ were well-explained by muscles spanning the hip during early stance – namely, the gluteus maximus and the biarticular hamstrings. We did not find a significant increase in metabolic cost at the hip during late stance when targeting smaller F_P_. However, at the individual muscle level, we did find higher metabolic costs of operating the psoas and iliacus during late stance. This outcome is consistent with increased demands for swing initiation, with metabolic consequences continuing into mid-swing, where we found a large increase in metabolic cost for all four quadriceps muscles. We interpret this increase to suggest that insufficient ankle power output exacts a metabolic penalty requiring additional effort for leg swing and knee extension in preparation for each subsequent foot strike. Together, these data provide an individual muscle roadmap to explain how the distal-to-proximal redistribution increases the metabolic cost of walking.

While walking with 40% smaller F_P_, we were surprised to find an increased metabolic cost of operating muscles spanning the ankle during single support. This increased utilization of ankle extensor muscles during single support may be due to a shift in the relative timing of the anterior GRF, which rose and peaked sooner for this biofeedback condition relative to normal walking (data not reported). At the individual muscle level, we were also surprised to find a reduced metabolic cost of operating the gluteus medius and minimus during push-off when targeting 40% smaller than normal F_P_. The gluteus medius and minimus are generally considered important during swing for regulating foot placement to preserve lateral stability. However, some authors have recently suggested that push-off intensity and lateral balance are inextricably connected (Kim & Collins 2015; Reimann et al. 2018). Our results are consistent with those conclusions and implicate the gluteus medius (and to a lesser extent, gluteus minimus) in providing hip stability that is proportional to push-off intensity, likely allowing for effective force transmission to the center of mass.

### Targeting larger than normal F_P_

The goal of many rehabilitation paradigms and wearable assistive devices is to restore power output during push-off. In the absence of such intervention, biofeedback that encourages increased push-off intensity shows promise to reverse the characteristic redistribution of joint power generation due to age (Lewis & Ferris 2008; Browne & Franz 2017a). Knowing the individual-muscle metabolic response to enhanced push-off intensity would allow scientists and clinicians to design better rehabilitation therapies for individualized prescription with translational potential to elicit more youthful and efficient gait biomechanics. Although our young adult population are not representative of those that would require such intervention, our data provide an important reference for future bioenergetic modeling studies in the target populations.

Our simulations reveal an increased metabolic cost of operating muscles spanning the hip, knee, and ankle when targeting larger than normal F_P_ – a response that manifested most during single support. At the hip, this metabolic penalty is explained primarily by increases for the gluteus medius and maximus during early single support and the hamstrings during midstance. Those changes may arise from larger than normal braking forces during early stance and larger acceleration of the center of mass during midstance, respectively. We predict that these metabolic changes would be mitigated if walking speed was permitted to increase in response to larger propulsive forces. This methodological design may also explain why we found no increase in the metabolic cost of muscles spanning the ankle during trailing limb double support when targeted larger than normal F_P_. Although this outcome is surprising, it is consistent with recent ultrasound imaging data alluding to a potentially counterproductive trade-off between increased activation and shorter operating lengths on force transmission from the calf muscles during push-off (Browne & Franz In Revision).

In contrast to effects when targeting 40% larger F_P_, and despite higher whole-body metabolic cost on average, targeting a more modest 20% increase in F_P_ resulted in lower metabolic costs to operate muscles spanning the hip and knee during leading limb double support. Here, the hamstring muscles appear to consume lower average metabolic cost during early stance compared to Norm. Thereafter, these reductions are overcome by increased costs during single support and leg swing. This suggests it may be theoretically possible to reduce metabolic costs of operating individual leg muscles to values lower than predicted for normal walking. Although at odds with generally accepted theories regarding energy optimization in walking (Browne & Franz 2017b), such a conclusion at the individual muscle level may allude to patterns of inter-muscular coordination that are outside the scope of this study but warrant further investigation.

Compared to walking with smaller F_P_, changes in the metabolic costs of operating individual leg muscles are much more evenly allocated when walking with larger F_P_. It is difficult to predict whether to expect the same outcome in older adults or people with gait pathology in response to interventions designed to restore F_P_. However, our results suggest that such interventions should consider not only the metabolic consequences of local (e.g., ankle) assistance to increase F_P_, but also any resultant compensation at other joints. Moving forward, we envision rehabilitative programs and assistive technologies that are informed by and objectively evaluated via individual muscle metabolic responses derived via the bioenergetic simulations used in this study.

### Limitations

We first acknowledge other factors that contribute to increased metabolic cost of walking in older adults. These may include changes in trunk flexion (Carey & Crompton 2005; Boyer et al. 2017), muscle biochemistry and mitochondria content (Brooks et al. 2005), and decreased tendon stiffness (Onambele et al. 2006). Our study instead used experimental manipulations in younger subjects to test for the metabolic consequences of biomechanical factors in the absence of other known age-related changes. We have reasonable evidence that the F_P_ biofeedback provides a method for young adults to emulate patterns of joint kinetics characteristic of older adults (DeVita & Hortobagyi 2000; Boyer et al. 2017; Browne & Franz 2017a; Browne & Franz 2019).

However, because our participants were not older adults, future work will need to test the sensitivity of these simulations to variations in model parameters. Another limitation we recognize is that we did not measure kinematics for the subtalar or metatarsal-phalangeal joints or the upper body. To overcome part of this limitation, we used trunk kinematics averaged from another cohort in our simulations. For transparency, our appendix documents these methodological considerations in more detail. Finally, participants walked at a fixed speed despite biofeedback-induced changes in F_P_. This led to differences in stride frequency known to affect metabolic cost (Donelan et al. 2002; Kuo et al. 2005; Soo & Donelan 2010) and likely to confound our results in ways described earlier.

## Conclusion

This study provides individual muscle-level insight into the metabolic consequences of walking with less and more vigorous push-off during walking. We first conclude that walking with insufficient push-off intensity places higher mechanical demand on muscles spanning the hip which in turn increases the metabolic cost of walking. This suggests that a distal-to-proximal redistribution of muscle demand during walking may independently contribute to higher metabolic costs in older adults and those with gait pathology. We also conclude that a fundamentally different and more evenly-allocated pattern of increased muscular demand explains higher metabolic energy costs when targeting larger propulsive forces. More generally, our simulations provide an initial benchmark for the metabolic consequences of altered propulsive force generation during walking that can inform strategies to improve walking economy in those with deficits in push-off intensity. Finding ways to preserve or restore ankle power generation, via rehabilitation, assistive technologies, or other means, appears central not only to improve walking performance, but also to improve walking economy.

## Acknowledgements

This study was supported by grants from the National Institutes of Health (R01AG058615) and Parkinson’s Foundation (PF-VSA-SFW-1908). We thank our participants for donating their time to participate in this study. We also thank Emily McCain, Rebecca Krupenevich, Gabriella Diaz, and Sidney Baudendistel for their expertise and assistance in study design, data collection, data processing, and modeling procedures.

## Appendix

### Model Efficacy

For the first 5 supplementary figures we show simulation results for all subjects, across all conditions, and on both sides for transparent reporting of simulation environments. In the final two supplementary figures we show results by condition averaged bilaterally and across participants.

In Supplementary Figure S1 we show joint kinematics which varied consistently across subjects, mainly occurring in the sagittal plane. In Supplementary Figure S2, we display the “hand of god” global forces and moments required to support each model in its environment while walking. The horizontal dashed black lines show the thresholds commonly recommended as optimal for a given trial (25 N forces & 75 Nm moments per OpenSim documentation). Some measurable deviations were present, tended to occur during double support, and were mostly concentrated in the vertical forces and sagittal plane moments. Joint moment reserves (i.e., those not accounted for by modeled muscle-tendon units) in Supplementary Figure S3 had fewer outliers than the global forces and moments, but a few persisted especially regarding hip flexion, hip adduction, hip rotation, and knee flexion.

Subject 10 was our primary outlier for reserve moments and forces (light orange), which we note carried through to the muscle forces (Supplementary Figure S4) and activation levels (Supplementary Figure S5). Specifically, we note that the hamstrings and gracillis muscle outcomes differed from other simulations between 40% and 60% of the gait cycle. We processed other strides for this participant and condition, and all yielded a similar response. We note that this one outlier occurred in the 40% larger F_P_ condition and may disproportionately contribute to the elevated metabolic cost of the biarticular hamstring muscles shown in Figure 4. Reassuringly, although the average profile for that condition was likely affected by this outlier, our SPM analysis correctly identified this region to have no condition main effect.

Moving on, muscle forces, normalized to body mass, were stable and, other than for Subject 10 as described, were fairly similar across all subjects (Supplementary Figure S4). Computed muscle activation levels (Supplementary Figure S5) were mostly uniform and temporally similar across all subjects. Though, Subject 2 tended to consistently activate muscles at a higher level than most other subjects. When averaging for each condition, the simulated muscle forces (Supplementary Figure S6) and activation levels (Supplementary Figure S7) displayed many features that align with the smaller and larger F_P_, similar to the metabolic cost profiles presented in our primary document.

### Addressing Model Efficacy

We understand that some of our simulations do not follow best practice recommendations (Hicks et al. 2015). The virtual trunk kinematics are likely the primary source of these discrepancies. The only other viable option to address the lack of torso kinematics was to place the full mass of the torso within the pelvis segment. However, we were concerned that this would affect the model center of mass to a higher degree than a virtual torso segment. Thus, we pressed on with the virtual torso kinematics superimposed on each participant’s lower body kinematics.

Another potential source of error may arise from scaling anatomical markers between the original model and their locations that depend on each participant’s anthropometrics. During the scaling process, we identified the average anatomical marker root mean square error of 0.016±0.004 m *(mean ± standard deviation),* with a max anatomical marker root mean square error of 0.0459 m *(Subject 3, left posterior superior iliac spine)*. We also note that subjects with higher BMIs tended to have higher scale error. Many of the anatomical marker locations help define limb segment dimensions and coordinate axes, thus errors during the model scaling process may carry forward into future model estimates.

Even with these limitations, we were able to demonstrate typical ranges of kinematics (even for the trunk), realistic muscle activation levels, and stable muscle forces across the gait cycle. To mitigate the effect of any simulation outliers, we averaged each pair of left and right-sided trials together when analyzing our primary outcome measure – muscle metabolic power. Finally, when reporting our metabolic costs, we present the individual data points for each gait cycle period of interest so that our readers may consider the results on an individual-subject level alongside the averages or boxplots for the group or condition. We believe this analysis can make a significant contribution to the field to understand individual muscle metabolic costs while walking across a range of F_P_, however we suggest that future investigations address the limitations we mention above.

## Supplemental Figure Captions

**Supplemental Figure S1:**
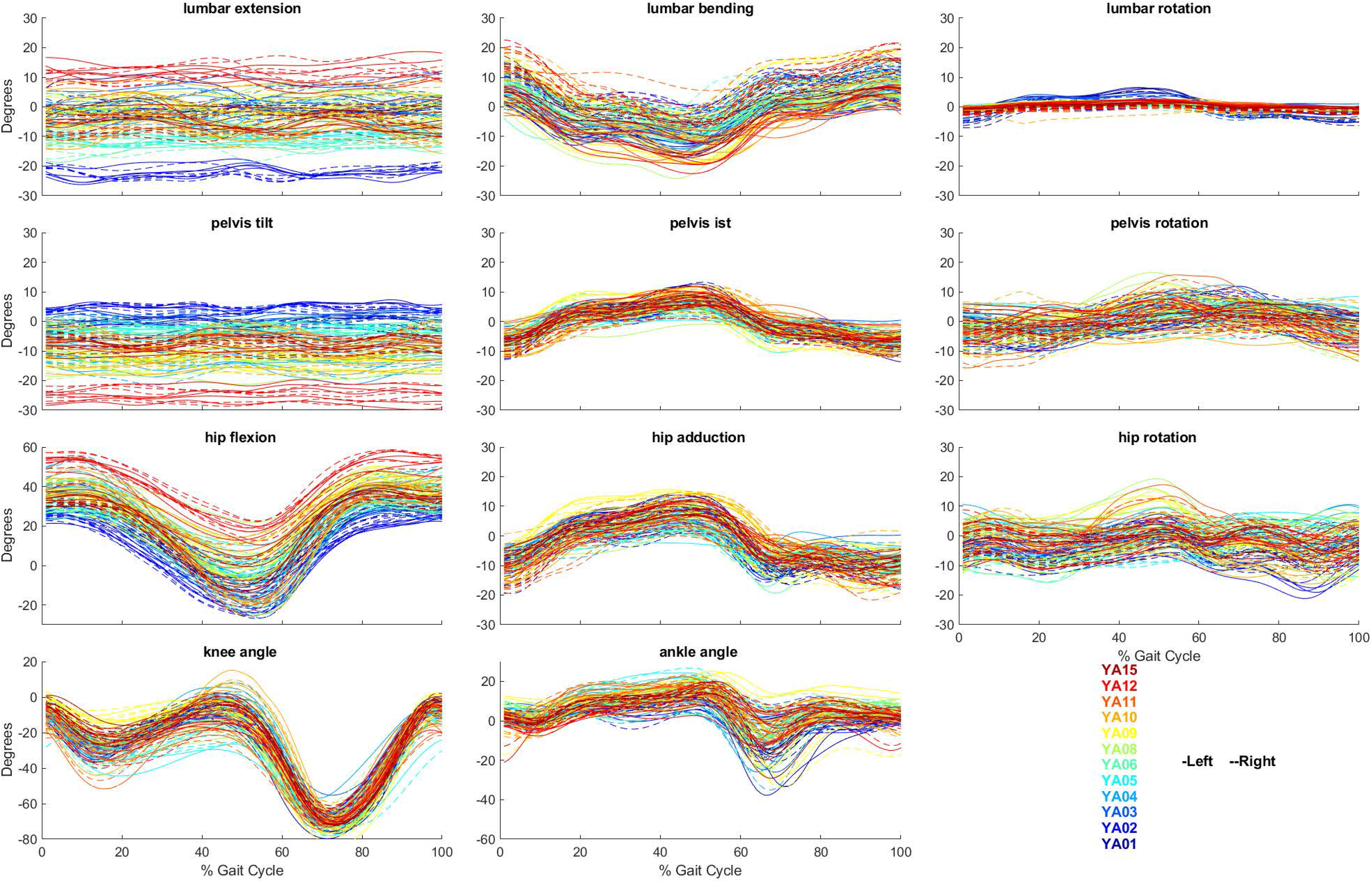
Joint kinematics across the gait cycle for all conditions in each participant.

**Supplemental Figure S2:**
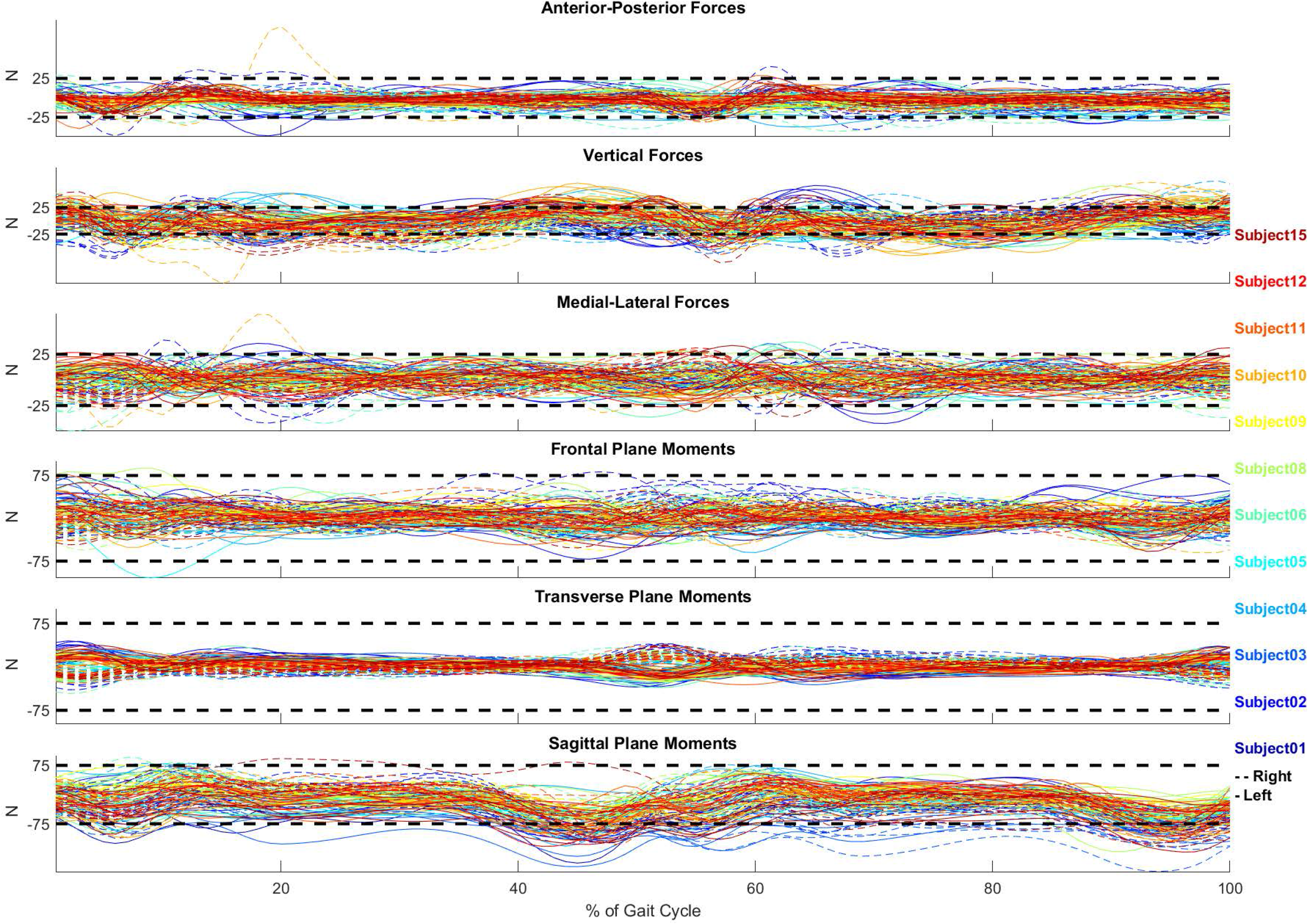
Global forces and moments across the gait cycle for all conditions in each participant.

**Supplemental Figure S3:**
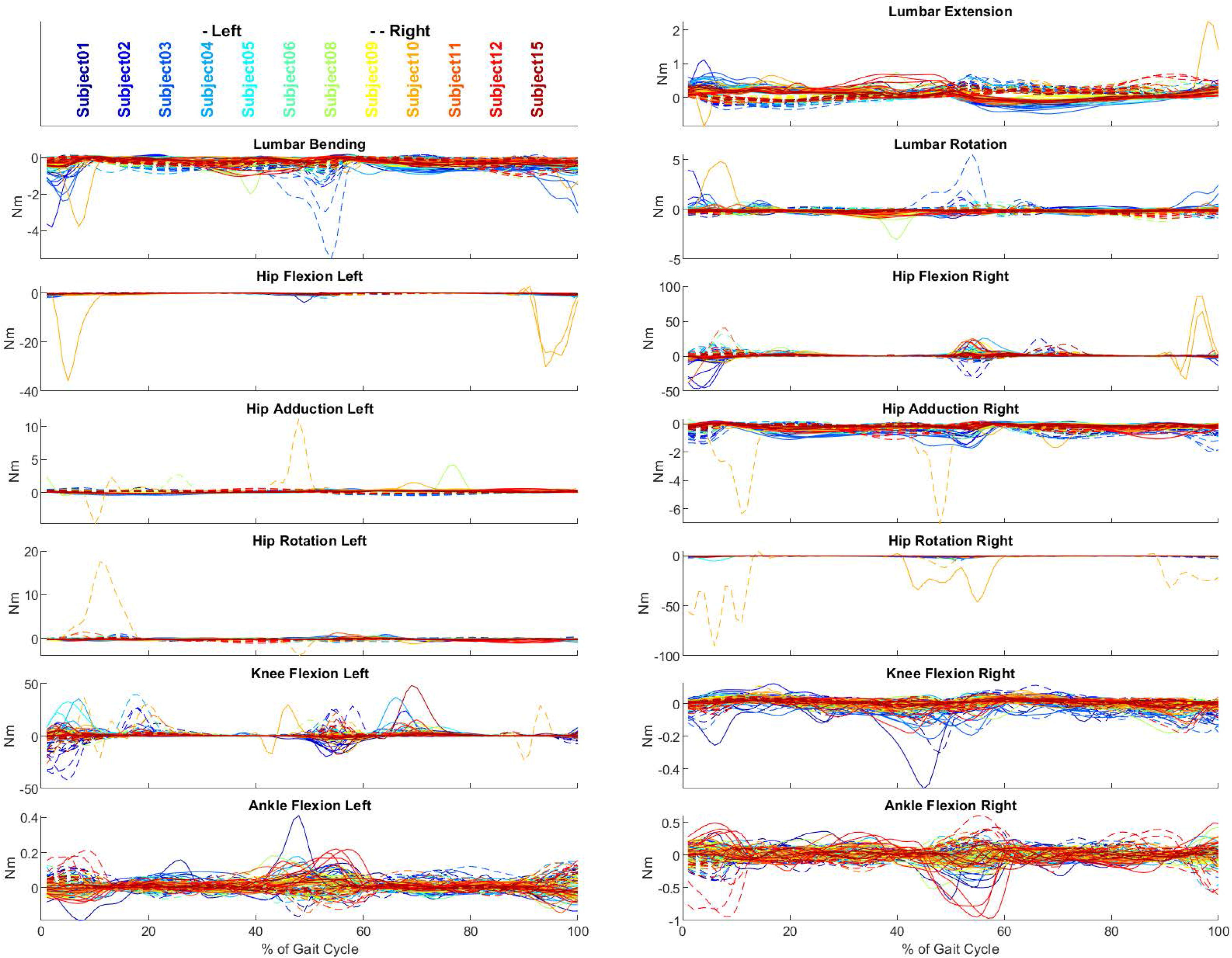
Net joint moment reserves moments across the gait cycle for all conditions in each participant.

**Supplemental Figure S4:**
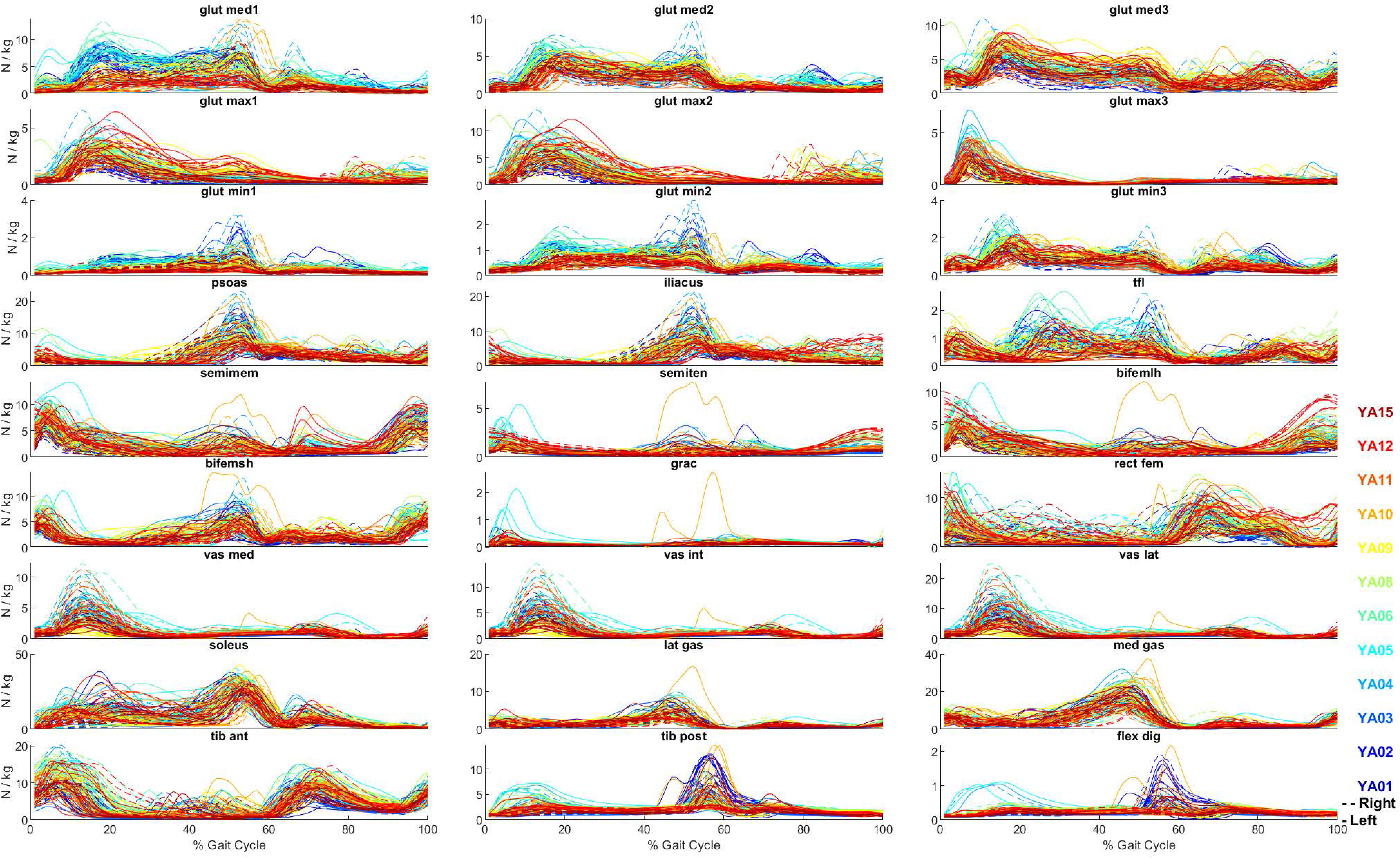
Computed muscle activation levels across the gait cycle for all conditions in each participant.

**Supplemental Figure S5:**
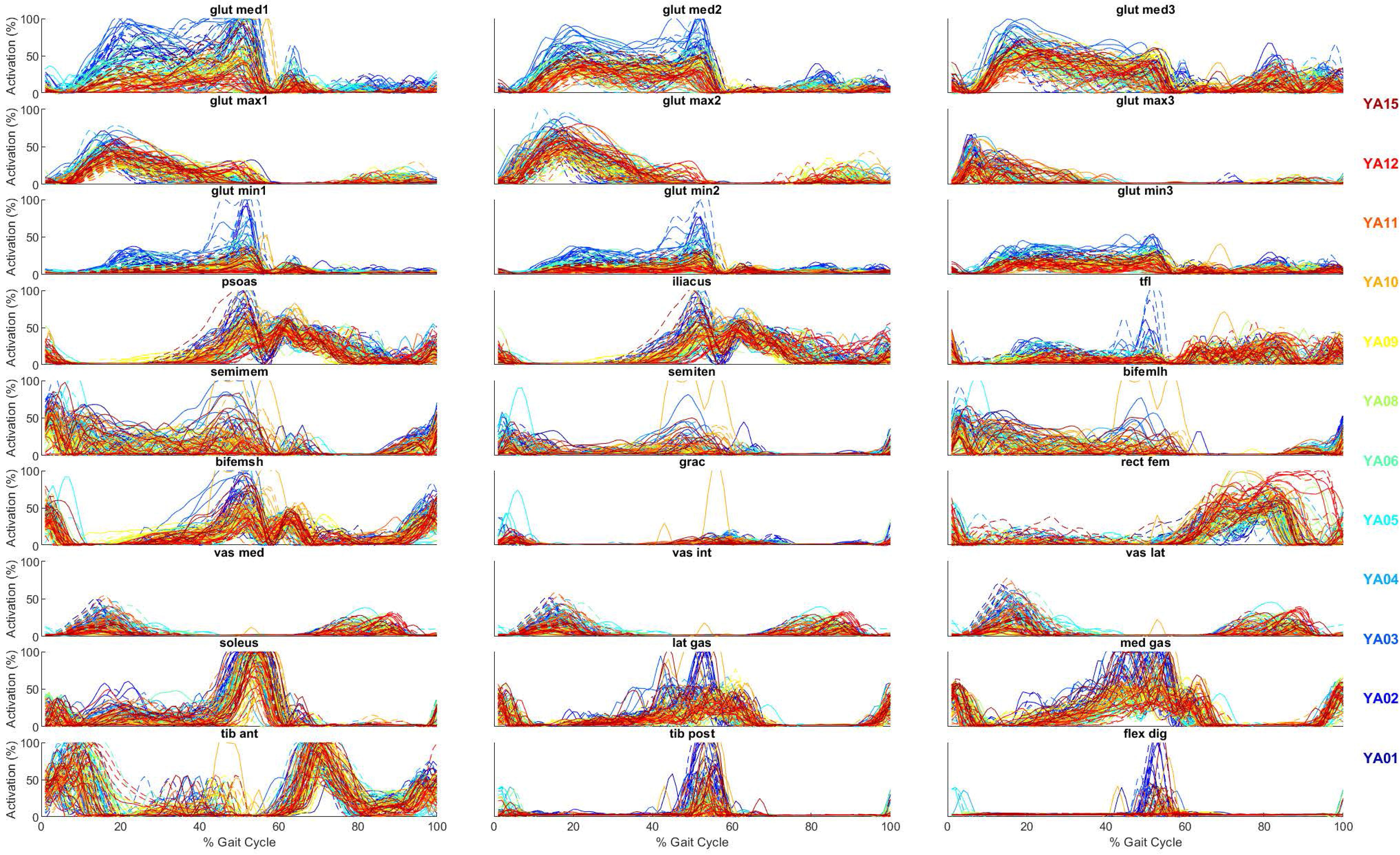
Relative simulated activation levels across the gait cycle for all conditions in each participant.

**Supplemental Figure S6:**
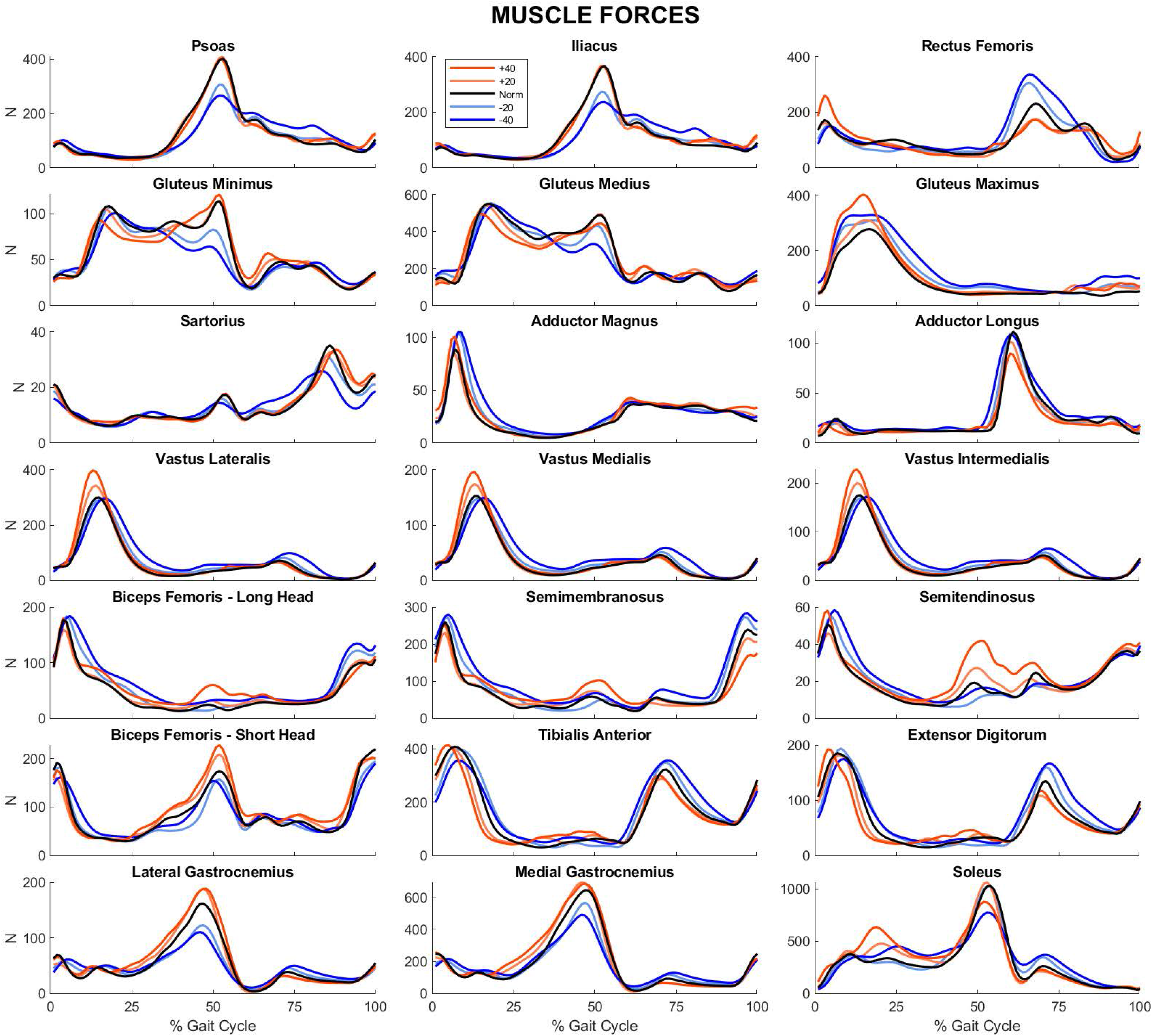
Condition average muscle forces for each of the 21 highest power-consuming muscles of the lower limb.

**Supplemental Figure S7:**
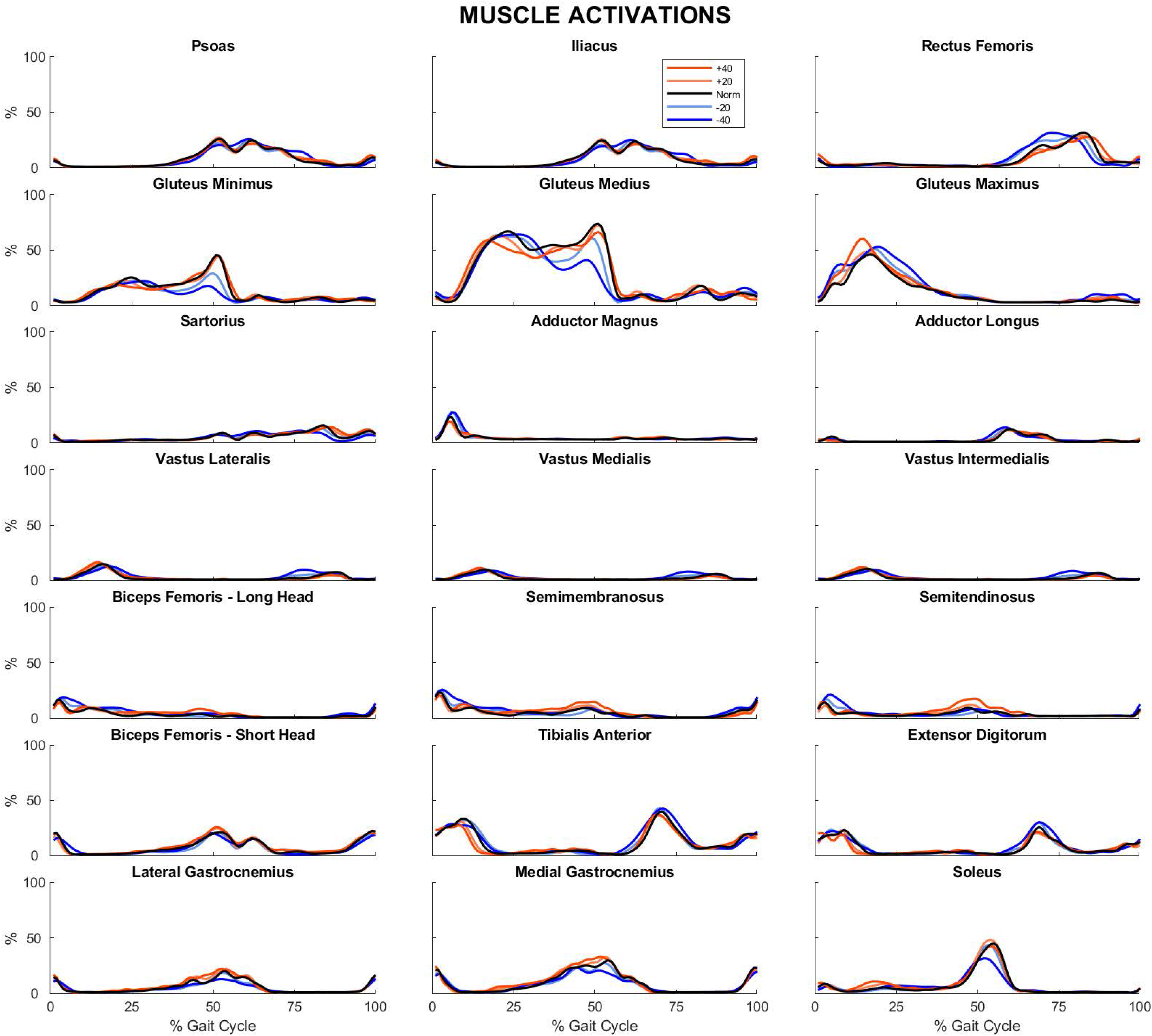
Condition average muscle activation levels for each of the 21 highest power-consuming muscles of the lower limb.

